# Sexually dimorphic leading-edge serrations evolved with sperm competition in silent swallows

**DOI:** 10.1101/2023.08.12.552953

**Authors:** Masaru Hasegawa

## Abstract

Leading-edge serrations are specialized feather structures, mitigating acoustic noise during foraging flight in owls, and have been extensively studied and applied to man-made noise-reducing structures. Similar structures occur in swallows, although the ecological functions of the serrations in these species remain unclear. I conducted macroevolutionary analyses of hirundines (Aves: Hirundininae), in which leading-edge serrations have evolved multiple times, to examine their evolution in relation to ecological context. I found that silent hirundines showed a higher probability of possessing leading-edge serrations, indicating that leading-edge serrations of swallows have an acoustic function as in owls. However, the probability of possessing leading-edge serrations had no detectable association with prey type in hirundines, indicating that their acoustic functions would evolve independent of foraging adaptation in this clade. Instead, the presence of leading-edge serrations was positively associated with relative testes mass, a well-known index of sperm competition, in hirundines. These findings explain sexually dimorphic expression of leading-edge serrations, a unique characteristic in this clade. In contrast to owls, in which serrations are considered an adaptation for silent foraging, sperm competition would favor the evolution of leading-edge serrations in hirundines, which illustrates that totally different selection pressures favor homologous traits in two taxa.

## INTRODUCTION

Leading-edge serrations, also known as leading-edge comb, are curved hooklets on the leading edge of the primary feathers in owls (reviewed in Wagner et al. 2017; Jaworski & Peake 2020; Terrill & Schultz 2023). This structure has intensively been studied and shown to reduce noise in flight and thus can be adaptation to improve hearing during foraging flights and to avoid detection by prey (e.g., Rao et al. 2017; reviewed in Wagner et al. 2017; Clark et al. 2020), which is now applied to man-made structure to reduce acoustic noise (e.g., pantograph of Japanese Shinkansen trains: reviewed in Wagner et al. 2017). However, owls are not the only animals that possess leading-edge serrations (Clark et al. 2020); similar structures, albeit somewhat smaller and less curved, can be found in other avian groups, including frogmouths (family: Podargidae) and swallows (family: Hirundinidae), and the function of the serrations in these birds remains unknown (reviewed in Clark et al. 2020).

In swallows (family: Hirundinidae), leading-edge serrations are known to have evolved independently at least twice, once in rough-winged swallows (genus: *Stelgidopteryx*; Fig. 1 left panel) and the other in saw-wings (genus: *Psalidoprocne*; Fig. 1 middle panel), evident from their English names (Turner & Rose 1994; also see Fig. 1 right panel for non-serrated leading-edge, typical to other hirundines). Similar to other species with leading-edge serrations (see above), these structures might serve to mitigate flight noise. However, different from the majority of owls and frogmouths, swallows are diurnal, visually-foraging birds, and, furthermore, they possess sexually dimorphic leading-edge serrations, in which males, but not females, possess serrations (reviewed in Turner & Rose 1994). This suggests that they might serve somewhat different functions than sexually monomorphic leading-edge serrations found in mainly nocturnal, acoustic predators (i.e., owls and frogmouths; e.g., see Roulin et al. 2013). In fact, although its function has not been shown, several hypotheses are proposed to explain the evolution of this sexually dimorphic structure (reviewed in Lunk 1962); for example, a clasping structure for copulation, to prevent slipping from females (Steinbacher 1931) and a means for generating non-vocal sounds during courtship or territorial chases (Lunk 1962; de Jong 2020). It is also unclear, if leading-edge serrations are adaptive, why only a few swallows possess leading-edge serrations (Steinbacher 1931; Lunk 1962), which sharply contrasts with owls in which virtually all owl species possess some forms of leading-edge serrations (e.g., Le Piane & Clark 2022).

**Figure 1.**
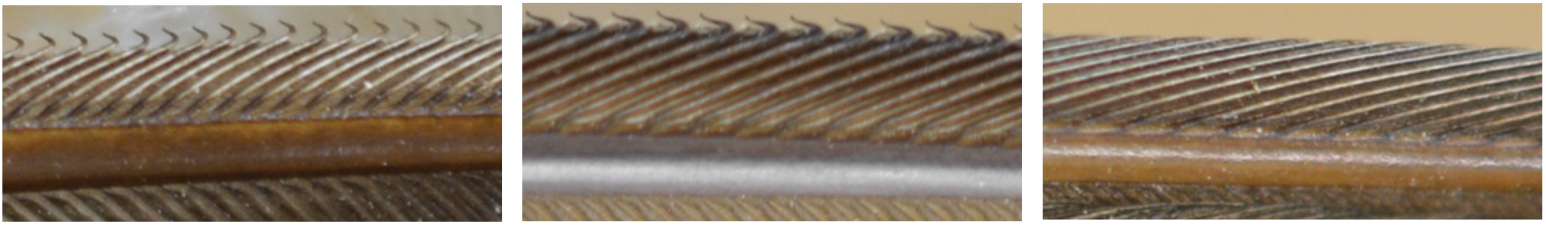
Examples of “leading-edge serrations” in hirundines: left panel) serrations of a male southern rough-winged swallow, *Stelgidopteryx ruficollis*; middle panel) serrations of a male white-headed saw-wing, *Psalidoprocne albiceps*, right panel) non-serrated leading-edge of a male red-rumped swallow, *Hirundo* (*Cecropis*) *daurica*.

Here, I studied ecological correlates of leading-edge serrations in swallows and martins (subfamily: Hirundininae). Because leading-edge serrations evolved at least two times (see above), phylogenetic comparative approaches can be used to find out ecological factors promoting the evolution of leading-edge serrations. Hirundines share a similar ecological background, such as aerial foraging, typically social monogamy, and biparental provisioning (Turner & Rose 1994), and thus would be suitable to conduct a macroevolutionary study focusing on ecological details (see below). Of several hypotheses proposed, the most promising hypothesis is that, as in owls, leading-edge serrations serve a function to reduce noise in flight (see above). Then, I predicted that vocal activity would be negatively linked to the evolution of leading-edge serrations. This is because reducing acoustic noise cannot make sense when birds continue to vocalize.

To further infer evolutionary forces favoring the leading-edge serrations, I also examined their association with foraging adaptations and sperm competition. Given that silence during foraging on elusive prey favors the leading-edge serrations in owls (see above), I expected that foraging on large and hence active fliers might favor the evolution of leading-edge serrations. This is because only active fliers can evade predation by detecting approaching predators. In other words, small, passive fliers (or, “aerial plankton”) cannot evade predation even if they perceive nearby predators. This foraging adaptation hypothesis is a possible explanation for the occurrence of leading-edge serrations in hirundines, but it does not account for their sexual dimorphism. Thus, I also examined leading-edge serrations in relation to the well-known measure of sperm competition, measured as testes mass (e.g., Møller 1991; Pitcher et al. 2005; Rowe et al. 2015). As secretary activity is favored in mating strategy as a “sneaker” (Shuster 2010), I predicted that leading-edge serrations would be evolutionarily positively associated with testes mass.

## MATERIAL & METHODS

### Data collection

As in previous studies (e.g., Hasegawa & Arai 2020a,b), information on morphological traits (i.e., bill size and relative tail fork depth, defined as fork depth divided by central tail feather length; e.g. Hasegawa & Arai, 2017, 2018) and migratory habits (i.e., migrant or not) was obtained from Hasegawa & Arai (2020b), which was derived from Turner & Rose (1994). Likewise, I collected data concerning the vocal activities (silent or not). Species with “silent” and “quiet” noted in the “voice” category was regarded as silent species (e.g., “they [southern rough-winged swallows] are generally silent”; “This [the black rough-winged swallow] is said to be a very silent swallow”; “Roughwings are generally silent”; “Brown-bellied swallows are usually quiet”; “they [Mascalene martins] are fairly quiet swallows”) and other species were regarded as non-silent species (unless data was unavailable). Dichotomous prey category (i.e., large and small flying insects), based on the size of prey item (roughly corresponding to >5mm or not), was obtained from Hasegawa & Arai (2022), which was originated from Turner & Rose (1994). The dichotomous prey category explains foraging traits (e.g., social/solitary foraging, foraging morphology; Hasegawa & Arai 2022; Hasegawa 2023a) and thus is a valid index of prey type in hirundines (see Hasegawa & Arai 2022; Hasegawa 2023a for details). In short, prey size is tightly linked to their flight speed, and large and small prey here roughly correspond to active fliers and aerial planktons, respectively (Hasegawa & Arai 2022; Hasegawa 2023a). Bill size, which did not impose a reduction of sample size in the current dataset, was used as another proxy of prey size (Fitzpatrick 1985; Hasegawa et al. 2016; Hasegawa & Arai 2023). Information on body mass, as a measure of body size, was obtained from Dunning (2008).

Information on testes mass as a measure of sperm competition, was obtained from a large dataset for avian species (Dun et al. 2001; also see Calhim & Birkhead 2007, in which the data set can be downloadable). Because this dataset does not contain information on testes mass for species with leading-edge serrations, I also searched for the information on testes mass for males in the breeding condition for these species (i.e., *Psalidoprocne* and *Stelgidopteryx*) using past publications and the information on museum specimens (*Psalidoprocne pristoptera,* n_male_ = 1: Mees 1970; *Stelgidopteryx ruficollis* n_male_ = 4, three from Rising 1961, one from collections of the Florida Museum of Natural History and *S. serripennis*, n_male_ = 2, one from Schaldach et al. 1997 and one from collections of the Florida Museum of Natural History). During this search for the information, I also obtained additional information on testes size of the cave swallow, *Petrochelidon fulva* (n_male_ = 6, four from Klass 1968 and two from collections of the Florida Museum of Natural History), the Bahama swallow, *Tachycineta cyaneoviridis* (n_male_ = 1, from collections of the Florida Museum of Natural History) and the blue swallow, *Hirundo atrocaerulea* (n_male_ = 1, Mees 1970), which were therefore included in the analysis (n_species_ = 18). To ensure compatibility across data sources, testes size measurements were converted to testes mass using the formula provided by Dun et al. (2001). To account for interspecific variation in body size, I used relative testes mass (i.e., testes mass/body mass) in statistical analyses. The complete dataset used is provided in Table S1. It was not possible to record data blind because presence/absence of serrations was evident from their English names (see above).

### Statistics

A Bayesian phylogenetic mixed model with a binomial error distribution was used to examine the presence/absence of leading-edge serrations. As in previous studies (e.g., Hasegawa & Arai 2020a,b; Hasegawa 2023b), potential confounding factors, body size [measured via log(body mass)] and migratory habit, were included alongside vocal activity in the model. I also included one of two indices of prey size (i.e., bill size and dichotomous prey size based on the size of prey items; see above) to test the association between prey size and the presence of leading-edge serrations. To account for phylogenetic uncertainty, I fit the models to each tree and applied a multimodel inference (e.g., Garamszegi & Mundry 2014), using 1000 alternative trees for swallows from birdtree.org (see Jetz et al. 2012, 2014; also see Sheldon et al. 2005 for an original phylogeny on swallows) with the prior [list(G = list(G1 = list(V = 1, nu = 200, alpha.mu = 0, alpha.V = 1)), R = list(V = 1, fix = 1))]. 1000 trees are sufficient to control for phylogenetic uncertainty (Rubolini et al. 2015). The residual variance was set at 1, in accordance with recommendations for categorical dependent variables (de Villemereuil et al. 2013). I used a Gelman prior for the fixed effects while standardizing each variable using “gelman.prior” in the package “MCMCglmm.” I derived the mean coefficients, 95% credible intervals (CIs) based on the highest posterior density, and Markov chain Monte Carlo (MCMC)-based P-values, together with the phylogenetic signal (de Villemereuil & Nakagawa 2014). Similarly, a Bayesian phylogenetic mixed model with a normal error distribution was used to examine relative testes mass in relation to the presence/absence of leading-edge serrations [prior: list(G = list(G1 = list(V = 1, nu = 200, alpha.mu = 0, alpha.V = 1)), R = list(V = 1, nu = 200))]. To better fit normal distribution in this analysis with a limited sample size, relative testes mass was log-transformed before analysis. In addition, relative tail fork depth, an intersexually selected trait in this clade (Hasegawa & Arai 2022), was included as a covariate. All analyses were conducted in R ver. 4.1.0 (R Core Team 2021) using the “MCMCglmm” function in the “MCMCglmm” package (ver. 2.29; Hadfield 2010). I ran each analysis for 140,000 iterations, with a burn-in period of 60,000 and a thinning interval of 80 for each tree.

A discrete module in BayesTraits (Pagel & Meade 2006) was used to examine evolutionary transitions among states with vocal activities (silent or not) and the presence/absence of leading-edge serrations. I ran for 1,010,000 iterations with a burn-in period of 10,000 and a thinning interval of 1000 (note that the phylogenetic tree was randomly chosen from 1000 trees in each iteration). I denoted means as the representatives of each transition rate. The reproducibility of the MCMC simulation was assessed using the Brooks–Gelman–Rubin statistic (Rhat), which must be <1.2 for all parameters (Kass et al. 1998), by repeating the analyses three times. Bayes factor was calculated via the stepping stone sampler in BayesTraits, comparing the marginal likelihood of a dependent model that assumed correlated evolution of the two traits (i.e., leading-edge serrations and vocal activity) to that of an independent model assuming independent evolution. Bayes factors of >2, 5–10, and >10 indicate positive, strong, and very strong evidence of correlated evolution, respectively (Kass & Raftery 1995; BayesTraits Manual Ver. 3).

## RESULTS

### Leading-edge serrations and silent vocal activity

Probability of possessing leading-edge serrations was significantly explained by the vocal activity of swallows (Table 1): after controlling for phylogeny, hirundines with relatively silent vocal activities showed a higher probability of possessing leading-edge serrations (Fig. 2). Three covariates, body size measured as log(body mass), migratory habits (migrant or not), and prey size measured as bill length were non-significant (Table 1, upper column). This was also the case when dichotomous prey size category (i.e., large vs small) was used instead of bill length (Table 1, lower column).

**Table 1.**
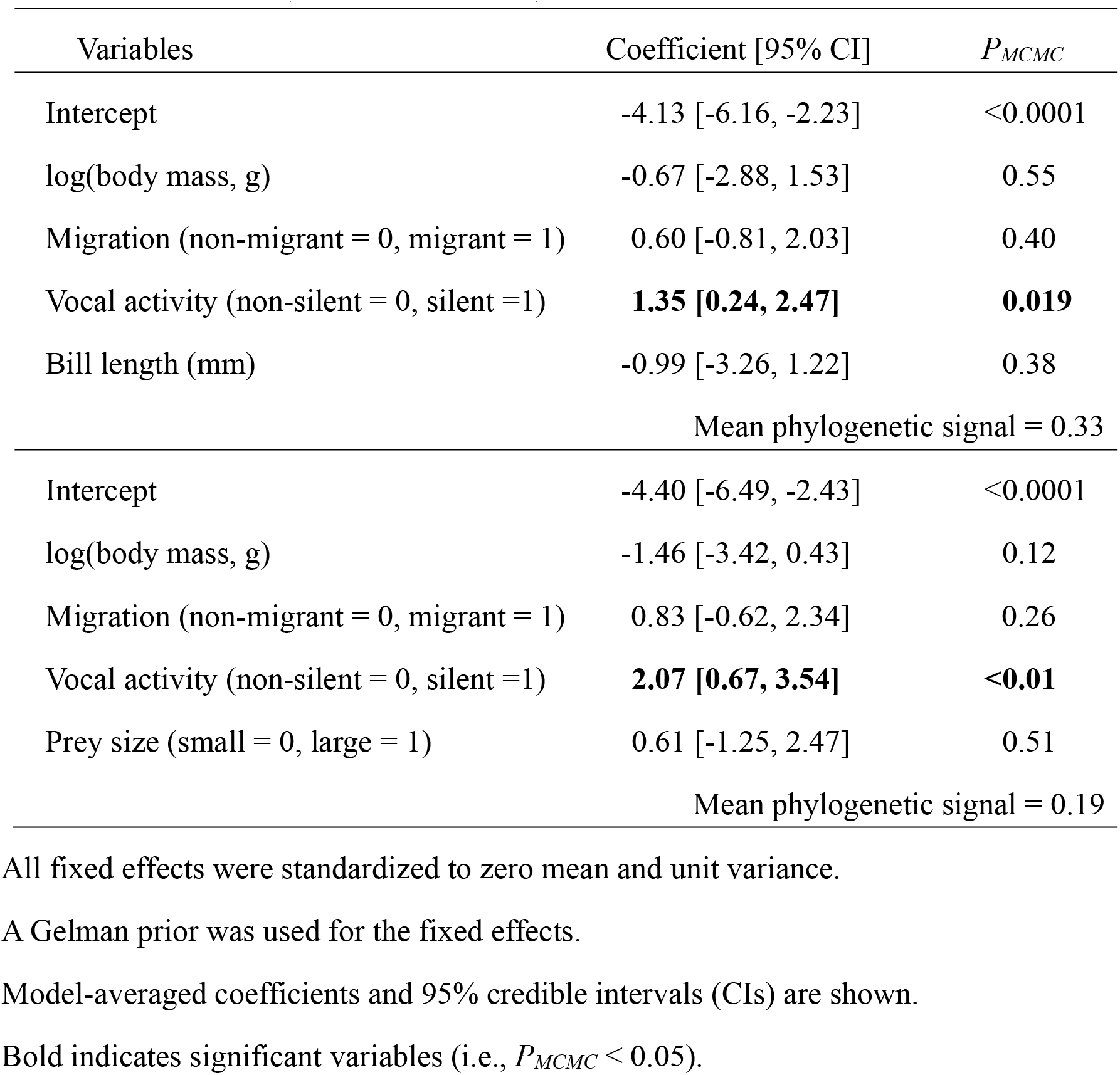
Multivariable Bayesian phylogenetic mixed model with a binary error distribution predicting the presence/absence of leading-edge serrations in relation to vocal activities and covariates including indices of prey size (upper column, bill length: n = 65; lower column, dichotomous prey size group, based on food items: n = 54) in swallows and martins (Aves: Hirundininae).

| Variables | Coefficient [95% CI] | $P_{MCMC}$ |
| --- | --- | --- |
| Intercept | -4.13 [-6.16, -2.23] | <0.0001 |
| log(body mass, g) | -0.67 [-2.88, 1.53] | 0.55 |
| Migration (non-migrant = 0, migrant = 1) | 0.60 [-0.81, 2.03] | 0.40 |
| Vocal activity (non-silent = 0, silent = 1) | **1.35 [0.24, 2.47]** | **0.019** |
| Bill length (mm) | -0.99 [-3.26, 1.22] | 0.38 |
| Mean phylogenetic signal = 0.33 |  |  |
| Intercept | -4.40 [-6.49, -2.43] | <0.0001 |
| log(body mass, g) | -1.46 [-3.42, 0.43] | 0.12 |
| Migration (non-migrant = 0, migrant = 1) | 0.83 [-0.62, 2.34] | 0.26 |
| Vocal activity (non-silent = 0, silent = 1) | **2.07 [0.67, 3.54]** | **<0.01** |
| Prey size (small = 0, large = 1) | 0.61 [-1.25, 2.47] | 0.51 |
| Mean phylogenetic signal = 0.19 |  |  |

All fixed effects were standardized to zero mean and unit variance.
A Gelman prior was used for the fixed effects.
Model-averaged coefficients and 95% credible intervals (CIs) are shown.
Bold indicates significant variables (i.e., $P_{MCMC} < 0.05$ ).

**Figure 2.**
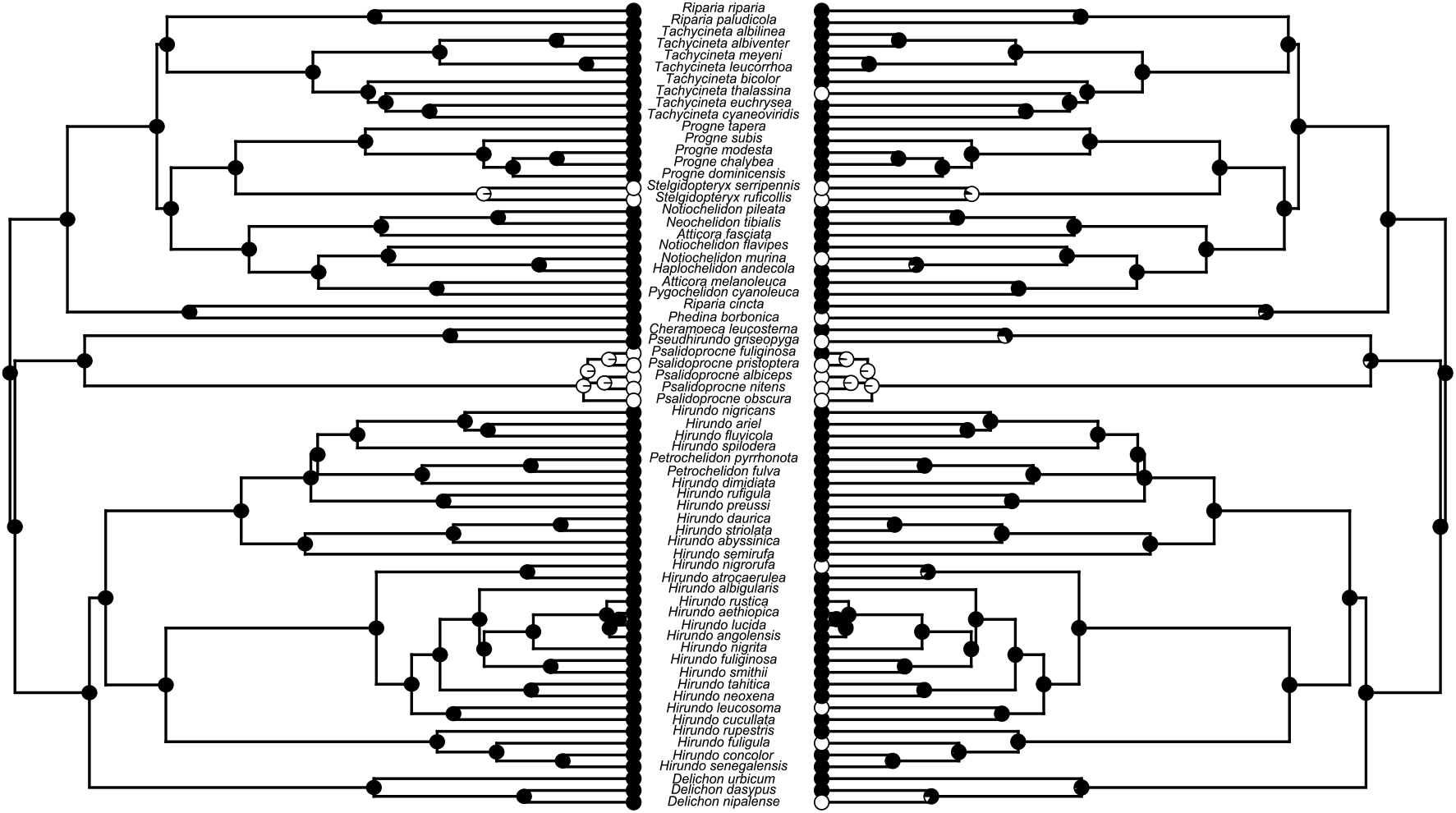
An example of ancestral character reconstruction of the leading-edge serrations (left) and of silent vocal activity (right) in swallows and martins (Aves: Hirundininae). Left side: presence and absence of leading-edge serrations are indicated by white and black circles, respectively; silent and non-silent vocal activity are indicated by white and black circles, respectively. The proportions of white and black at the nodes indicate the probability of the ancestral state. Here, I used the functions “ace” in the R package “ape” (with model = “ER”, i.e., an equal-rates model; Paradis et al. 2005) and “plotTree” in the R package “phytools” (Revell 2012) for ancestral character reconstruction.

When examining evolutionary transitions among states (i.e., presence/absence of leading-edge serrations and silent/non-silent vocal activities), I found very strong support for correlated evolution between these two state variables (n = 68, Bayes Factor = 12.43). Transition rate to the state with leading-edge serrations was lower than the reverse transition in silent hirundines (Mean difference = -57.65, P_MCMC_ = 0.014), which was not the case in non-silent hirundines (Mean difference = -13.10, P_MCMC_ = 0.59; Fig. 3), though the results should be carefully treated as leading-edge serrations evolved only a few times, most likely, two times (Fig. 2).

**Figure 3.**
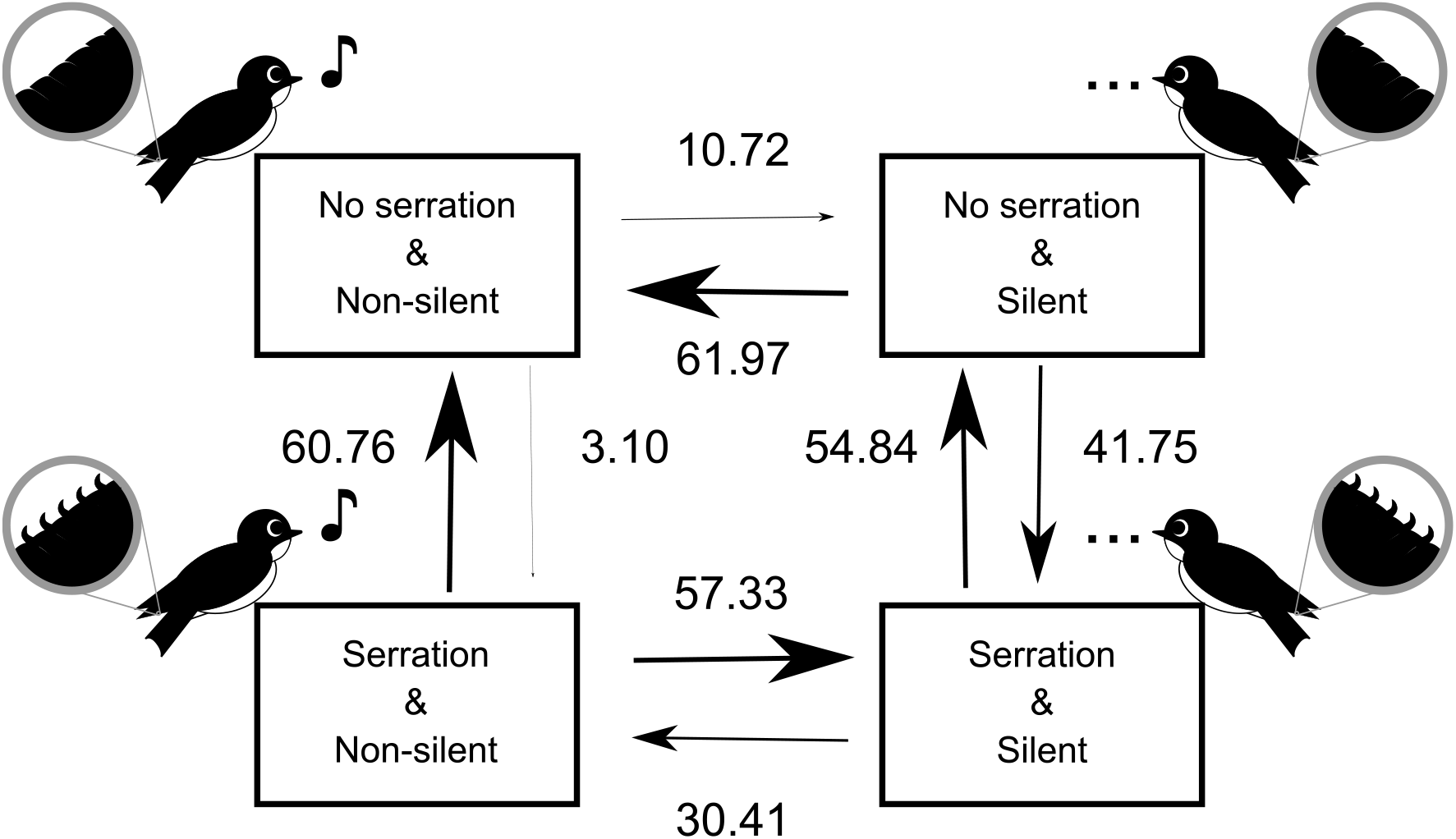
The most likely evolutionary transitions between states with the presence/absence of leading-edge serrations and the silent/non-silent vocal activity (see illustrations beside each state) in swallows and martins (Aves: Hirundininae). Numbers next to arrows, represented by arrow size, indicate transition rates between pairs of states.

### Leading-edge serrations and sperm competition

Testes mass relative to body mass was positively associated with the presence of leading-edge serrations when controlling for relative tail fork depth as an intersexually selected trait (Table 2; Fig. 4). This relationship remained significant even when relative tail fork depth was excluded from the model (univariable Bayesian phylogenetic mixed model: Coefficient = 1.48, 95% CI = 0.13, 2.86, P_MCMC_ = 0.033).

**Table 2.** Multivariable Bayesian phylogenetic mixed model with a normal error distribution predicting relative testes mass, log(testes mass/body mas), in relation to the presence/absence of leading-edge serrations and relative tail length as an intersexually selected trait (n = 18) in swallows and martins (Aves: Hirundininae).

| Variables | Coefficient [95% CI] | $P_{MCMC}$ |
| --- | --- | --- |
| Intercept | -0.29 [-0.92, 0.34] | 0.34 |
| Relative tail length (mm) | 0.39 [-0.14, 0.92] | 0.14 |
| Leading-edge serrations<br>(absence = 0, presence = 1) | **1.66 [0.27, 3.04]** | **0.02** |
| Mean phylogenetic signal = 0.12 |  |  |

Relative tail length was standardized to zero mean and unit variance. Model-averaged coefficients and 95% credible intervals (CIs) are shown. Bold indicates significant variables (i.e., $P_{MCMC} < 0.05$ ).

**Figure 4.**
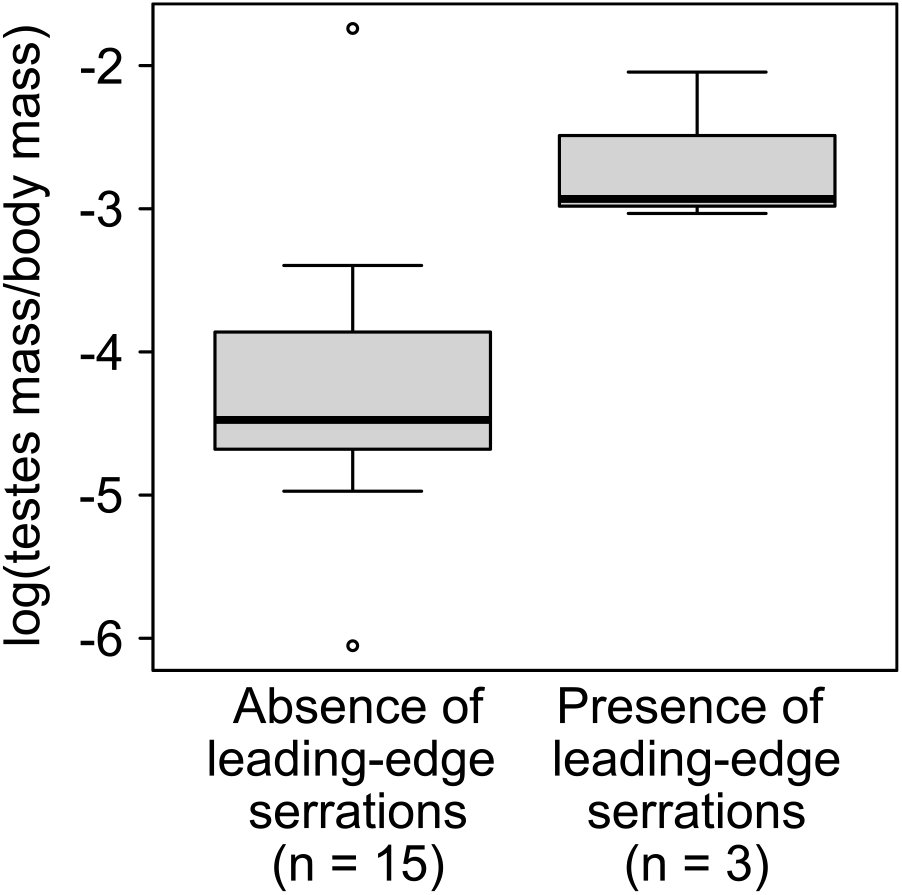
Boxplots of log(testes mass/body mass) for species with and without leading-edge serrations in hirundines. The horizontal bar in each boxplot indicates the median, and the box shows the first and third quartiles of data. The whiskers range from the lowest to the highest data points within 1.5 × interquartile range of the lower and upper quartiles, respectively (see Table 2 for the formal statistical analysis).

## DISCUSSION

The main finding of the current study is that leading-edge serrations evolved from relatively silent hirundines, which cannot be explained by the possibility that serrations evolved and are maintained as a clasping structure for copulation (see Introduction). Instead, the current study supports that leading-edge serrations have an acoustic function. Silent vocal activity evolved in a limited number of branches, mostly in relatively young ones (Fig. 2), explaining the limited occurrence of leading-edge serrations in hirundines. Although the current study does not clarify the exact acoustic function, they may reduce noise in flight as found in owls and in artificial wing models (e.g., Rao et al. 2017; see Introduction).

I also found that relative testes mass, a proxy of sperm competition (see Introduction), was positively associated with the presence of leading-edge serrations, indicating that these serrations would coevolve with intense sperm competition. In acoustic-signaling animals, selection can favor signal loss (or, silence) through the modification or reduction of sound-producing structures (e.g., Hawaiian crickets; Pascoal et al. 2014). Likewise, selection for silent male swallows might favor deformed leading-edges. For example, leading-edge serrations (and their likely consequence, silent flight) could aid in extra-pair mating by reducing detection by rival males or by decreasing the likelihood of female evasion, thereby enhancing extra-pair and within-pair mating success (e.g., see Hasselquist & Bensch 1991 for an example of “silent sneaking” in birds). In fact, even without leading-edge serrations, flight noise is inaudible from a distance (Wagner et al. 2017), and thus leading-edge serrations would be functional only in a close-range, as in mating interaction.

An alternative possibility that hirundines use leading-edge serrations to reduce noise during foraging flight (see Introduction) would be unlikely, as leading-edge serrations had no detectable association with prey size (note that small, aerial planktons cannot evade predation even if they detect nearby predators and thus this hypothesis predicts that leading-edge serrations would not evolve in hirundines foraging on small prey items). In addition, this hypothesis alone cannot explain sexually dimorphic expression of leading-edge serrations. It should also be noted that serrations are thought to mitigate flight noise primarily during the terminal phase of prey capture, just before landing, in owls (Wagner et al 2017; Clark et al. 2017). This functional context does not readily apply to continuous aerial foraging in hirundines (Turner & Rose 1994). Taken together, these indicate that the overall pattern in hirundines is not consistent with the foraging adaptation hypothesis.

Another explanation of the observed pattern is that leading-edge serrations are used for non-vocal sound production during courtship or territorial chases (Lunk 1962; de Jong 2020). Silent vocal activity might be a pre-condition for the evolution of non-vocal acoustic communication. Although this is a fascinating idea, there is no evidence that leading-edge serrations can produce non-vocal sounds and thus is an unlikely explanation on the basis of a large amount of evidence that serrations reduce, rather than increase, acoustic noise (see Introduction). It is, however, possible that leading-edge serrations are a device to reduce noise to increase the detectability of seemingly non-vocal, subtle sounds reported in swallows with leading-edge serrations (i.e., genus: *Stelgidopteryx* and *Psalidoprocne*: Turner & Rose 1994; de Jong 2020), though we still need to identify the mechanisms of the possible non-vocal sounds.

In summary, I demonstrated here that leading-edge serrations co-evolved with silent vocal activities. The current study is the first macroevolutionary analyses of sexually dimorphic leading-edge serrations. Because the presence of leading-edge serrations was positively associated with the measure of sperm competition (but not with measures of prey type), intense sperm competition rather than foraging adaptation would explain sexually dimorphic expression of leading-edge serrations. In hyperaerial insectivores such as swallows, potential aerodynamic costs associated with leading-edge serrations (sensu Le Piane & Clark 2022; see also Rao et al. 2017 for reduced lift-to-drag ratios in serrated wings) might prevent females from evolutionarily maintaining male-like serrations, which is likely the case but should be tested in future. Although actual ecological function of leading-edge serrations in hirundines remains to be demonstrated using manipulative experiments (e.g., through removal of serrations), the observed pattern contrasts with that in owls and other nocturnal, acoustic birds, where sexually monomorphic leading-edge serrations likely confer a viability advantage by enhancing prey capture. The current study exemplifies that totally different selection pressures (i.e., foraging adaptation in owls and sperm competition in hirundines, here) can favor homologous traits (i.e., leading-edge serrations) in two taxa.

## ACKNOWLEDGMENTDS

I thank Dr Emi Arai, Dr Shumpei Kitamura and his lab members at Ishikawa Prefectural University for their kindest advices. I am grateful to Dr Angela Turner for her kindly support on the valuable information on swallows. I also thank Dr Glaucia Del-Rio and her lab members at Florida Museum of Natural History for their kindest support.

## DATA AVAILABILITY STATEMENT

The data sets supporting this article are included as Table S1.

## ETHICS

This comparative study does not include any treatments of animals, as all the information was gathered from bodies of literature.

---

**Table S1.** Dataset of the current study (n = 65).

| Scientific Name | Presence of<br>serrations | Body<br>mass (g) | Presence of<br>Migration | Silent<br>vocalization<br>(1) or not (0) | Bill<br>length<br>(mm) | Prey size<br>(large = 1) | Relative tail<br>fork depth | Testes<br>mass (g) |
| --- | --- | --- | --- | --- | --- | --- | --- | --- |
| *Neochelidon tibialis* | 0 | 9 | 0 | 0 | 8 | 0 | 0.28 | NA |
| *Stelgidopteryx ruficollis* | 1 | 15 | 1 | 1 | 9 | 1 | 0.00 | 1.94 |
| *Stelgidopteryx*<br>*serripennis* | 1 | 15 | 1 | 1 | 10 | 0 | 0.00 | 0.80 |
| *Tachycineta bicolor* | 0 | 20 | 1 | 0 | 9 | 0 | 0.17 | 0.67 |
| *Tachycineta albilinea* | 0 | 13 | 0 | 0 | 11 | 1 | 0.14 | NA |
| *Tachycineta albiventer* | 0 | 15 | 1 | 0 | 11 | 1 | 0.21 | NA |
| *Tachycineta thalassina* | 0 | 15 | 1 | 1 | 9 | 0 | 0.2 | NA |
| *Tachycineta leucorrhoa* | 0 | 19 | 1 | 0 | 11 | 1 | 0.13 | NA |
| *Tachycineta meyeni* | 0 | 17 | 1 | 0 | 10 | 1 | 0.16 | NA |
| *Tachycineta*<br>*cyaneoviridis* | 0 | 17 | 0 | 0 | 10 | 0 | 0.66 | 0.04 |
| *Notiochelidon murina* | 0 | 11 | 0 | 1 | 9 | NA | 0.41 | NA |
| *Notiochelidon flavipes* | 0 | 10 | 0 | 0 | 9 | NA | 0.29 | NA |
| *Notiochelidon pileata* | 0 | 12 | 0 | 0 | 7 | NA | 0.43 | NA |
| *Pygochelidon cyanoleuca* | 0 | 10 | 1 | 0 | 7 | 0 | 0.26 | 0.20 |
| *Atticora fasciata* | 0 | 14 | 0 | 0 | 8 | 0 | 0.89 | NA |
| *Atticora melanoleuca* | 0 | 11 | 0 | 0 | 7 | 0 | 0.97 | NA |
| *Progne tapera* | 0 | 36 | 1 | 0 | 15 | 1 | 0.26 | 0.41 |
| *Progne subis* | 0 | 56 | 1 | 0 | 14 | 1 | 0.30 | 0.48 |
| *Progne chalybea* | 0 | 41 | 1 | 0 | 14 | 1 | 0.31 | 0.99 |
| *Progne dominicensis* | 0 | 40 | 1 | 0 | 14 | 1 | 0.33 | NA |
| *Riparia paludicola* | 0 | 13 | 0 | 0 | 8 | 0 | 0.00 | NA |
| *Riparia riparia* | 0 | 13 | 1 | 0 | 9 | 1 | 0.21 | 0.12 |
| *Riparia cincta* | 0 | 21 | 1 | 0 | 12 | 1 | 0.00 | NA |
| *Psalidoprocne fuliginosa* | 1 | 11 | 0 | 0 | 7 | NA | 0.43 | NA |
| *Psalidoprocne albiceps* | 1 | 11 | 1 | 1 | 7 | 0 | 0.47 | NA |
| *Psalidoprocne*<br>*pristoptera* | 1 | 11 | 1 | 1 | 7 | 0 | 0.62 | 0.53 |
| *Psalidoprocne obscura* | 1 | 9 | 0 | 1 | 7 | 0 | 1.85 | NA |
| *Psalidoprocne nitens* | 1 | 9 | 0 | 1 | 7 | 0 | 0.00 | NA |
| *Cheramoeca leucosterna* | 0 | 14 | 0 | 0 | 8 | 0 | 1.06 | NA |
| *Pseudhirundo griseopyga* | 0 | 9 | 0 | 1 | 8 | 0 | 1.00 | NA |
| *Phedina borbonica* | 0 | 21 | 0 | 1 | 11 | 0 | 0.17 | NA |
| *Hirundo rupestris* | 0 | 23 | 1 | 0 | 11 | 1 | 0.00 | NA |
| *Hirundo fuligula* | 0 | 22 | 0 | 1 | 10 | 0 | 0.00 | NA |
| *Hirundo concolor* | 0 | 13 | 0 | 0 | 8 | NA | 0.00 | NA |
| *Hirundo rustica* | 0 | 18 | 1 | 0 | 12 | 1 | 1.45 | 0.31 |
| *Hirundo lucida* | 0 | 13 | 0 | 0 | 10 | 0 | 0.53 | NA |
| *Hirundo angolensis* | 0 | 17 | 0 | 0 | 11 | 0 | 0.35 | NA |
| *Hirundo tahitica* | 0 | 13 | 0 | 0 | 11 | 1 | 0.18 | 0.09 |
| *Hirundo neoxena* | 0 | 14 | 0 | 0 | 9 | 1 | 0.67 | 0.15 |
| *Hirundo albigularis* | 0 | 21 | 1 | 0 | 12 | 1 | 0.73 | NA |
| *Hirundo aethiopica* | 0 | 13 | 0 | 0 | 10 | 0 | 0.75 | NA |
| *Hirundo smithii* | 0 | 13 | 1 | 0 | 9 | 0 | 1.19 | NA |
| *Hirundo nigrita* | 0 | 17 | 0 | 0 | 11 | 1 | 0.14 | NA |
| *Hirundo dimidiata* | 0 | 11 | 1 | 0 | 9 | NA | 0.62 | NA |
| *Hirundo atrocaerulea* | 0 | 13 | 1 | 0 | 10 | 0 | 2.14 | 2.28 |
| *Hirundo nigrorufa* | 0 | 13 | 1 | 1 | 9 | 0 | 0.36 | NA |
| *Hirundo cucullata* | 0 | 27 | 1 | 0 | 10 | 0 | 1.00 | NA |
| *Hirundo abyssinica* | 0 | 17 | 1 | 0 | 9 | 0 | 1.00 | NA |
| *Hirundo semirufa* | 0 | 30 | 1 | 0 | 11 | 0 | 1.57 | NA |
| *Hirundo senegalensis* | 0 | 42 | 0 | 0 | 13 | NA | 1.10 | NA |
| *Hirundo daurica* | 0 | 19 | 1 | 0 | 11 | 0 | 1.40 | NA |
| *Hirundo striolata* | 0 | 22 | 0 | 0 | 11 | 1 | 1.09 | NA |
| *Hirundo preussi* | 0 | 13 | 0 | 0 | 8 | 0 | 0.26 | NA |
| *Hirundo rufigula* | 0 | 15 | 1 | 0 | 8 | 0 | 0.22 | NA |
| *Hirundo spilodera* | 0 | 20 | 1 | 0 | 8 | 1 | 0.00 | NA |
| *Hirundo fuliginosa* | 0 | 12 | 0 | 0 | 8 | NA | 0.13 | NA |
| *Petrochelidon*<br>*pyrrhonota* | 0 | 21 | 1 | 0 | 9 | 0 | 0.00 | 0.45 |
| *Haplochelidon andecola* | 0 | 17 | 0 | 0 | 9 | NA | 0.00 | NA |
| *Petrochelidon fulva* | 0 | 15 | 1 | 0 | 8 | 0 | 0.00 | 0.31 |
| *Hirundo fluviicola* | 0 | 9 | 1 | 0 | 8 | NA | 0.00 | NA |
| *Hirundo ariel* | 0 | 11 | 1 | 0 | 7 | NA | 0.15 | NA |
| *Hirundo nigricans* | 0 | 15 | 1 | 0 | 9 | 1 | 0.16 | 0.14 |
| *Delichon urbicum* | 0 | 18 | 1 | 0 | 9 | 0 | 0.50 | 0.20 |
| *Delichon dasypus* | 0 | 18 | 1 | 0 | 8 | 0 | 0.12 | NA |
| *Delichon nipalense* | 0 | 15 | 0 | 1 | 7 | 0 | 0.00 | NA |

NA indicates data “Not Available.”
See text for detailed information.

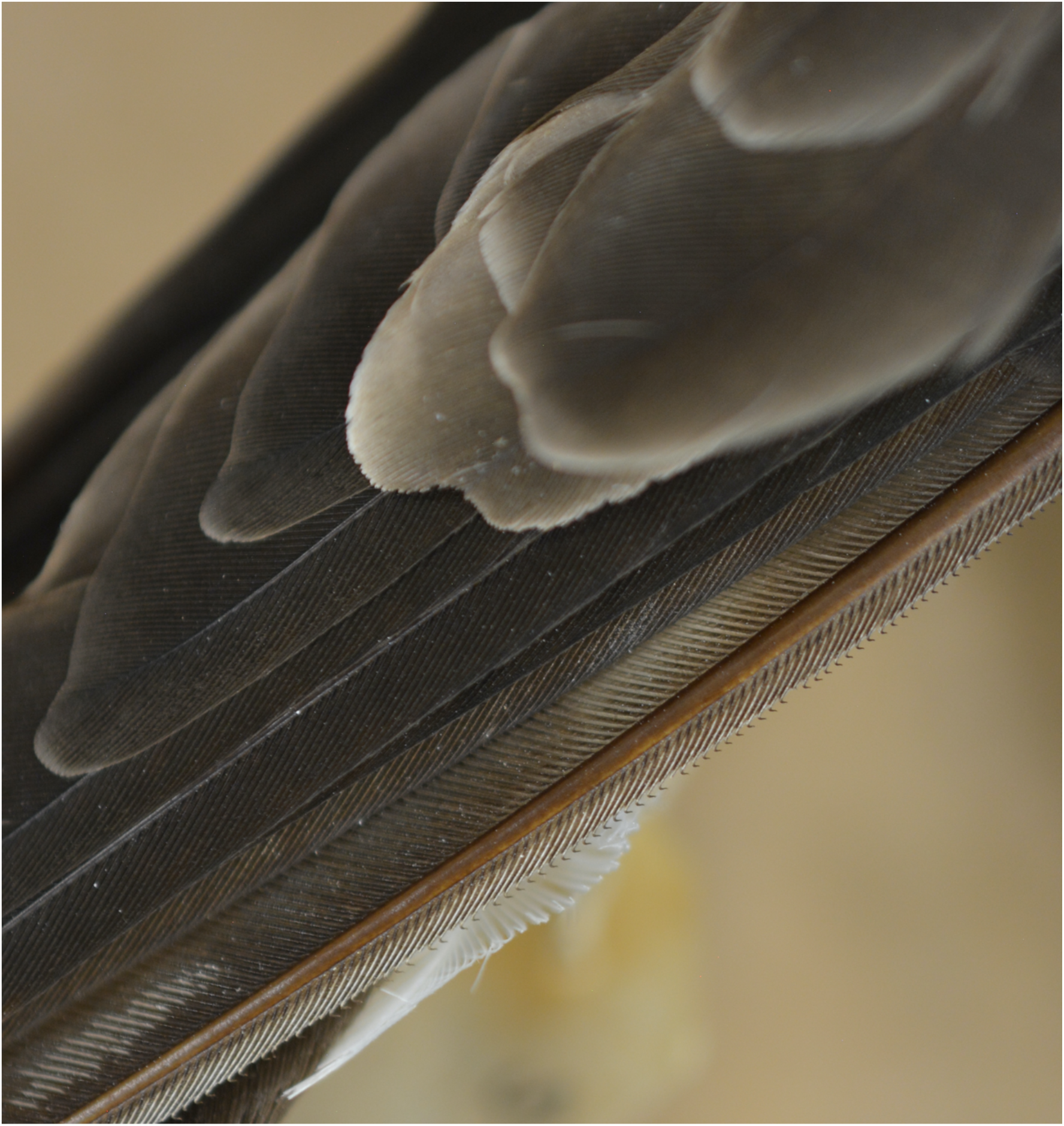
Cover image

